# Dietary carotenoid supplementation facilitates egg laying in a wild passerine

**DOI:** 10.1101/792234

**Authors:** Jorge García-Campa, Wendt Müller, Sonia González-Braojos, Emilio García-Juárez, Judith Morales

## Abstract

During egg laying, female birds face a trade-off between self-maintenance and investment into current reproduction. Providing eggs with resources is energetically demanding, since in most species females lay one egg per day. However, the costs of egg laying not only relate to energetic requirements, but also depend on the availability of specific resources that are vital for egg production and embryonic development. One of these compounds are carotenoids, pigments with antioxidant properties and immuno-stimulatory functions, which are crucial during embryonic development. In this study, we explore how carotenoid availability alleviates this trade-off and facilitates egg laying in the blue tit. Blue tit females lay one egg per day and have the largest clutch size of all European passerines. We performed a lutein supplementation experiment, and measured potential consequences for egg laying capacity and egg quality. We found that lutein-supplemented females had less laying interruptions and thus completed their clutch faster than control females. No effects of treatment were found on the onset of egg laying or clutch size. Experimentally enhanced carotenoid availability did not elevate yolk carotenoid levels or egg mass, but negatively affected eggshell thickness. Our results provide hence evidence on the limiting role of carotenoids during egg laying, However, the benefits of laying faster following lutein supplementation were counterbalanced by a lower accumulation of calcium in the eggshell. Thus, even though single components may constrain egg laying, it is the combined availability of a range of different resources which ultimately determines egg quality and thus embryonic development.

## Introduction

Life-history theory predicts that increased investment into current reproduction provides immediate fitness benefits via enhanced reproductive success, while it impinges at the same time on the amount of resources that can be maintained for self-maintenance and thus for future reproduction (Stearns 1992). In birds, females face this trade-off between current and future reproduction among others when allocating resources to their eggs, as this increases offspring viability, but the costs of egg production compromise their rearing capacity and their prospects for future reproduction as well as survival (Monaghan et al. 1998; Visser and Lessells 2001). The high costs of egg production and the difficulties to maintain egg quality throughout laying are also reflected in the changes in egg composition along the egg sequence (Nager et al. 2000; Williams and Miller 2003).

Variation in egg composition along the laying sequence relates on the one hand to the energetic requirements for egg production that involve the acquisition of nutrients to be allocated to the eggs (Carey 1996; Monaghan and Nager 1997). Such energy or nutrient related effects on egg production have been studied in food supplementation experiments (Harrison et al. 2010). Indeed, providing females with more nutrients advanced the timing of reproduction (Vafidis et al. 2016), and had positive effects on clutch size (Korpimäki and Hakkarainen 1991) or egg size (Ardia et al. 2006). Laying capacity depends on the other hand on the availability of specific resources that are essential for embryonic development. One of these essential dietary micronutrients are carotenoids. These pigments are involved in a wide range of physiological processes, playing among others an immuno-stimulatory (Pérez-Rodríguez et al. 2008) and antioxidant role (Cohen and McGraw 2009; Watson et al. 2018). Carotenoids are crucial at early stages of development, since they reduce embryonic ROS damage (Surai and Speake 1998) and enhance offspring immune system before (Surai et al. 2001) and after hatchling (McWhinney et al. 1989; Haq et al. 1996). Furthermore, they have an effect on offspring growth (Biard et al. 2007) and influence the development of traits like plumage or beak colouration that play an important role in parent-offspring communication (Tschirren et al. 2003; Morales and Velando 2013).

However, carotenoids cannot be endogenously synthetized by vertebrates and must be acquired from the diet. This implies that carotenoids could become limiting during highly demanding periods, and that their use could be constrained by both their availability or by an individual’s ability to find them. Thus, allocating carotenoids to eggs is expected to impose a physiological cost for females (Surai et al. 2001; McGraw et al. 2005; Karadas et al. 2005). Carotenoid demand for self-maintenance processes is greater during the breeding season and, particularly, during egg laying, a period framed by a situation of high oxidative stress and immune-depression (Sheldon and Verhulst 1996; Hansell et al. 2005). This challenging period may –depending on clutch size-be extensive, since carotenoid acquisition and transfer to the eggs starts already days prior to egg laying (e.g., in birds, 5 days prior to laying; Surai 2001). Egg quality is hence likely reduced under low carotenoid availability (Bortolotti et al. 2003). Indeed, previous studies have found that eggs laid by carotenoid supplemented females contained higher yolk carotenoid concentrations (see Blount et al. 2002; Biard et al. 2007).

There is also some evidence that egg production per se could be limited by low carotenoid availability (Blount et al. 2000, 2004), potentially with negative effects on clutch size in conditions of low carotenoid availability (Eeva and Lehikoinen 2010). Furthermore, when carotenoids are limited females may also extend the laying period by lowering their egg laying rate. In birds, females normally lay one egg per day until the clutch is completed, but laying interruptions of one or several days can occur (reviewed in Astheimer 1985; see also Nilsson and Svensson 1993). These interruptions have been reported to be more frequent under harsh conditions such as cold weather (Lessells et al. 2002), high pollution (Eeva and Lehikoinnen 2010) and poor calcium availability (Graveland 1996; Bureš and Weidinger 2003; Eeva and Lehikoinen 2010). Yet, such delays in the reproductive schedule may negatively affect fitness because they increase the time in which females and their clutches are vulnerable to predators (Milonoff 1989), may decouple the timing for chick rearing with the peak of food availability (Durant et al. 2005), or may increase hatching asynchrony, if females start incubating before clutch completion (Magrath 1990). However, no study has explored the effects of carotenoid availability on egg laying interruptions.

In this study, we explored the previous question in a small passerine, the blue tit (*Cyanistes caeruleus*). In this species, females make a substantial investment into their clutch, which can weigh up to 150% of their own body mass (Perrins and Birkhead 1983; Stenning 2018). As income breeders, blue tit females must acquire all the resources allocated to the clutch from their diet, for a period of up to 2 weeks. Dietary carotenoid availability is likely of central importance as it is known to have significant implications for blue tit embryonic and post-hatching development (e.g. Surai and Speake 1998; Biard et al. 2007; Valcu et al. 2019). Moreover, blue tit females supplemented with carotenoids at laying have been found to allocate more carotenoids to the egg yolk and to raise chicks with enhanced carotenoid-based coloration (Biard et al. 2006). We tested whether experimentally enhanced carotenoid availability prior and during laying affected the occurrence of laying interruptions, as well as clutch size and laying date. We predicted that carotenoid supplemented females would have less laying interruptions, may advance egg laying and lay larger clutches. We also explored whether carotenoid supplementation influenced various aspects of egg quality like the amount of carotenoids in the yolk, as well as egg mass and shell thickness.

## Material and methods

### General methods

The study was carried out in Miraflores de la Sierra, Community of Madrid, central Spain (40° 481⍰N, 03° 471⍰W) during the spring of 2017. We studied a nest-box breeding blue tit population in a deciduous forest that is dominated by Pyrenean oak (*Quercus pyrenaica*). The blue tit is a territorial-monogamous passerine that has one of the largest clutch size and the largest variation in clutch size among all European passerines (Stenning 2018). Eggs weigh on average 1.17 g (n= 1001; Stenning 2018), ranging from 0.97-1.41 g (Nilsson and Svensson 1993). As many other passerines (Perrins 1970), blue tit females lay one egg per day, but laying interruptions are frequent and are more abundant under environmental constraints and food unavailability (Nilsson and Svensson 1993; Matthysen et al. 2010; Stenning 2018).

From the beginning of April onwards, nest-boxes were visited every two days to determine the start of nest building. Blue tit nests are mainly made of moss, which is formed and lined with soft material such as hair and feathers. Nest building is mainly done by the female. As soon as the nest cup was defined (a hole in the moss not coated with soft material), which is typically 6.2 days prior to egg laying (range 1-14 days), we started lutein supplementation (see the following section).

Once egg laying started, nests were visited every second day and eggs were marked on the day they were found. Blue tit females lay one egg per day, although laying interruptions are frequent, so visiting nests every second day is sufficient to notice any laying interruptions. The fifth egg was collected immediately and substituted by a fake egg to prevent females from replacing it. Lutein supplementation (see below) continued throughout egg laying and stopped with the onset of incubation. Lutein supplementation lasted on average 15.7 days prior to clutch completion (range: 9-23 days), as blue tit females tend to start incubating before the clutch is complete (Salvador 2012; Stenning 2018). A total of 92 nests were included in this experiment, and the average clutch size was 9.36 (n=92, range 6-14).

We trapped adults during parental feeding visits (8-12 days after hatching; hatching date = day 0) by means of nest-box traps. Adults were weighed with a Pesola spring balance to the nearest 0.01 g.

In this study, we did not include hatching date effects due to a second experimental design in which we cross-fostered clutches two days before the expected hatching date. Thus, we avoided including hatching date in the analyses of the present study due to possible late prenatal effects related to incubation of foster females.

### Manipulation of lutein availability

Once the nest cup was defined but not lined, we put a transparent plastic feeder into the nest-box (2.5×4.5×4.5 cm), pinned to the inner back nest-box wall. We chose this stage of nest building to ensure that the nest owners would continue breeding, since most nest usurpations occur at earlier stages. Nests were sequentially assigned to either a control group or to a lutein-supplemented group. Initially, we created 60 control nests and 32 lutein-supplemented nests. This unbalance between treatments was designed in the context of the second experiment mentioned above, in which we needed twice the number of control nests than lutein-supplemented nests. In the present study, we nevertheless used all data available. One nest was deserted during food supplementation as result of a nest usurpation by another blue tit pair.

Lutein was provided every second day, and each dosage consisted of 50 mg of Versele Laga Yel-lux Oropharma (lutein 8000 mg/Kg), which corresponds to 0.4 mg of lutein. According to Partali et al. (1987), one lepidopteran larvae in the natural diet of blue tits contains on average 5.3 µg of lutein and thus the dosage used would correspond, approximately, to 75 prey items. At least, blue tits chicks consume on average 100 lepidopteran larvae per day (Gibb and Betts 1963), and thus our experimental manipulation every two days lies within the natural range. Each lutein dose was mixed with 5 g of commercial fat with nuts (GRANA Oryx). Control nests received the same amount of fat but without lutein. Each lutein dose was weighted in advance with a digital analytical balance (accuracy 0.001 mg) and stored in Eppendorf tubes at 4°C in the dark to prevent oxidation. Lutein doses were mixed with the corresponding amount of fat just before supplementation at the nest.

We supplemented nests until the female started incubation. At each visit, we weighed the amount of food that remained at the feeder using a Pesola spring balance (to the nearest 0.01 g), cleaned the feeder and refilled it with 5 g with the corresponding treatment. We then calculated the total amount of food that was consumed over the two days. We also noted the first day when at least 0.5 g of food was consumed, in order to estimate the number of days it took a given female to start consuming food (hereafter also termed “food neophobia”). In a subsample of nests (n=15), we confirmed by means of video recordings that males rarely visited the nest-box and, thus, we assume that any food that disappeared was consumed mainly by the female.

### Egg measurements

We collected the 5^th^ egg on the day of laying (n=80 clutches; 51 from control nests and 29 from lutein-supplemented nests) and weighed it in the field to the nearest 0.01 g. We registered egg mass in 79 eggs because one egg of the lutein-supplemented group was broken during the egg collection. Eggs were kept cool and within the same day they were stored at - 80°C until the analyses. For the analysis, we defrosted all eggs on the same day and separated the yolk, the albumen and the eggshell, and weighed the yolk using a digital analytical balance (accuracy 0.001 g). We added to the yolk twice the volume of water and vortexed this mixture at the highest speed for 1 min. Yolks were again stored at −80°C until the following day, when they were defrosted again for carotenoid analyses. We could use 76 samples for yolk mass and carotenoid analysis (48 from control nests and 28 from lutein-supplemented nests) because yolk and albumen was mixed in a few samples after defrosting.

From that yolk-water mixture, 100 μl were transferred to a new Eppendorf and 400 μl of pure ethanol were added. The mixture was then centrifuged at 1500 G during 5 min at room temperature, and the aqueous phase was transferred to an Eppendorf tube. Optical density was obtained at 450 nm using a Synergy™ HT Multi-model Microplate Reader (BioTek^®^ Instruments, Inc.). Carotenoid concentrations were obtained from a lutein analytical standard (Sigma-Aldrich^®^). Plate number was registered for each sample and controlled for in statistical analyses. In a subsample of eggs (n=30), we also analysed lutein concentration by high-performance liquid chromatography (HPLC) with ethanol extraction following Alonso-Álvarez et al. (2004), and both measurements were positively correlated (*r*_30_=0.41, *P*=0.02). We calculated yolk carotenoid content by multiplying carotenoid concentration and yolk mass.

Eggshell thickness of all the 80 collected eggs was measured using a digital tube micrometer (Mitutoyo Ip-65) with ball-point ends and precision of 0.001 mm, following Morales et al. (2013). We took 9 measures per egg (if possible, 3 in each of the following eggshell locations: blunt end, sharp end and equator). When it was not possible to identify the specific location of the eggshell, it was categorized as “indeterminate”. Measures at these different locations showed high repeatability (blunt end: *r*=0.8, *F* =10.7, *P* < 0.001; sharp end: *r*=0.8, *F* =12.7, *P* < 0.001; equator: *r*=0.8, *F* =17.3, *P* < 0.001; indeterminate: *r* =0.8, *F* =4.4, *P* < 0.001; all measures pooled: *r*=0.7, *F*_1,79_ =21.4, *P* < 0.001). Thus, we calculated the mean of each eggshell location. We clearly identified equator location for 48 shells, 33 shells for sharp end location and 24 for shells from blunt end location.

### Statistical Analyses

First, we tested whether the total amount of food consumed and food neophobia differed between the control and the lutein-supplemented group. Food neophobia was analysed using a Generalized Lineal Model (GLZ) with Poisson distribution and a log link function, including treatment, the total amount of food consumed, female body mass and all double interactions with treatment as predictor variables. The total amount of food consumed was analysed with General Linear Models (GLM), including treatment, female body mass and the interactions between both as predictor variables.

Second, we tested the effect of treatment on laying capacity variables (laying date, number of laying interruptions and clutch size) and egg quality (egg mass and yolk carotenoid content) by using GLMs, except for the number of laying interruptions, which was analysed using a GLZ with Poisson distribution. In these models, we included treatment, the total amount of food consumed, female body mass and all double interactions between treatment and the rest of the parameters as predictor variables. We also included clutch size and its interaction with treatment as predictor variables in the analysis of laying interruptions. Plate number was included as predictor in yolk carotenoid analysis.

Finally, shell thickness was analysed using a mixed model including nest id as random factor in order to account for repeated measures at different shell locations, and treatment, the total amount of food consumed, female body mass, eggshell location (blunt end, sharp end or equator) and all double interactions with treatment as fixed predictor variables.

We used SAS 9.4 (SAS Inst., Cary, NC, USA) for all statistical analyses. Backward elimination of non-significant interactions (α=0.05) was used to acquire minimal models. The models were checked for residual normality with a Shapiro normality test. In the text, we report minimal models after backward elimination, while full models are shown in tables.

## Results

### Total amount of food and food neophobia

Treatment did not significantly affect the total amount of food consumed (coef. = −3.86 ± 1.85, *F*_1,69_ = 3.77, *P* = 0.052), although lutein-supplemented females tended to eat more food than control females. Lutein-supplemented females showed less food neophobia than control females (coef. = 0.39 ± 0.18, *χ*^2^_1_ = 4.87, *P* = 0.027). Food neophobia did not depend on female’s body mass or the total amount of food consumed (both *P* > 0.1). None of the interactions with treatment was significant (Table 1).

**Table 1.**
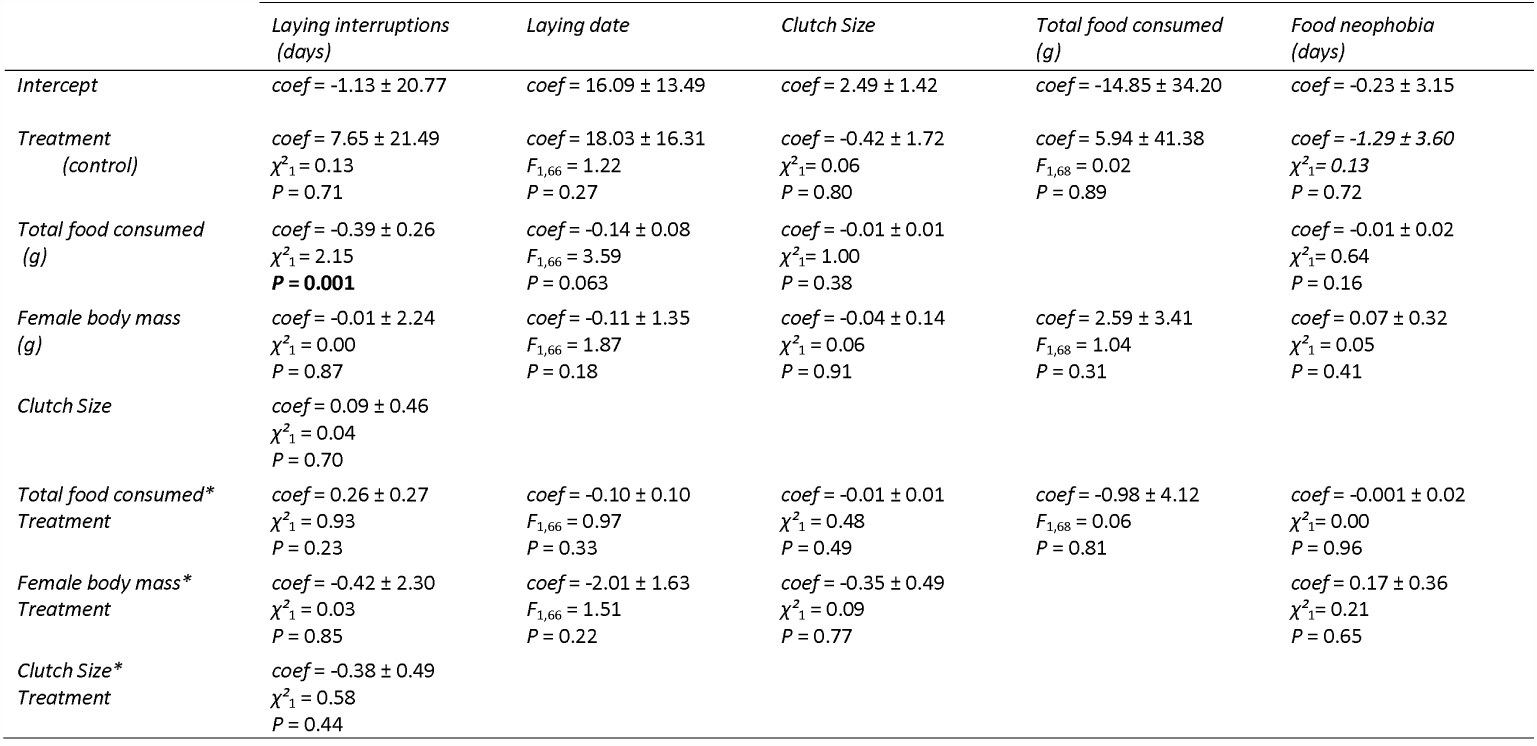
Full models before backward deletion of non-significant interactions showing the effect of carotenoid supplementation on laying capacity and food consumption. General lineal models were performed for laying date and food neophobia analyses. Laying interruptions and clutch size models were performed using Generalized lineal models. Coefficients are shown for control nests. Significant differences are marked in bold.

### Egg laying capacity

There was no treatment effect on the onset of egg laying (coef. = −1.20 ± 0.80, *F*_1,68_ = 2.28, *P* = 0.14) or on clutch size (coef. = 0.03 ± 0.08, *χ*^2^_1_ = 0.1, *P* = 0.75). Lutein-supplemented females had less laying interruptions than control females (coef. = 1.17 ± 0.63, *χ*^2^_1_ = 4.54, *P* = 0.033). Females that consumed more food had less laying interruptions (coef. = −0.14 ± 0.05, *χ*^2^_1_ = 11.30, *P* < 0.001), while no effects of female body mass were found (coef. = −0.35 ± 0.49, *χ*^2^_1_ = 0.54, *P* = 0.46). All interactions with treatment were not significant (see Table 1).

### Egg quality

No effects of treatment (coef. = 0.02 ± 0.02, *F*_1,59_ = 0.75, *P* = 0.39) or of the total amount of food consumed (coef. = −0.001 ± 0.001, *F*_1,59_ = 1.32, *P* = 0.25) were found on egg mass. Heavier females laid heavier eggs (coef. = 0.06 ± 0.02, *F*_1,59_ = 11.92, *P* = 0.001). The interactions with treatment were not significant (Table 2).

**Table 2.**
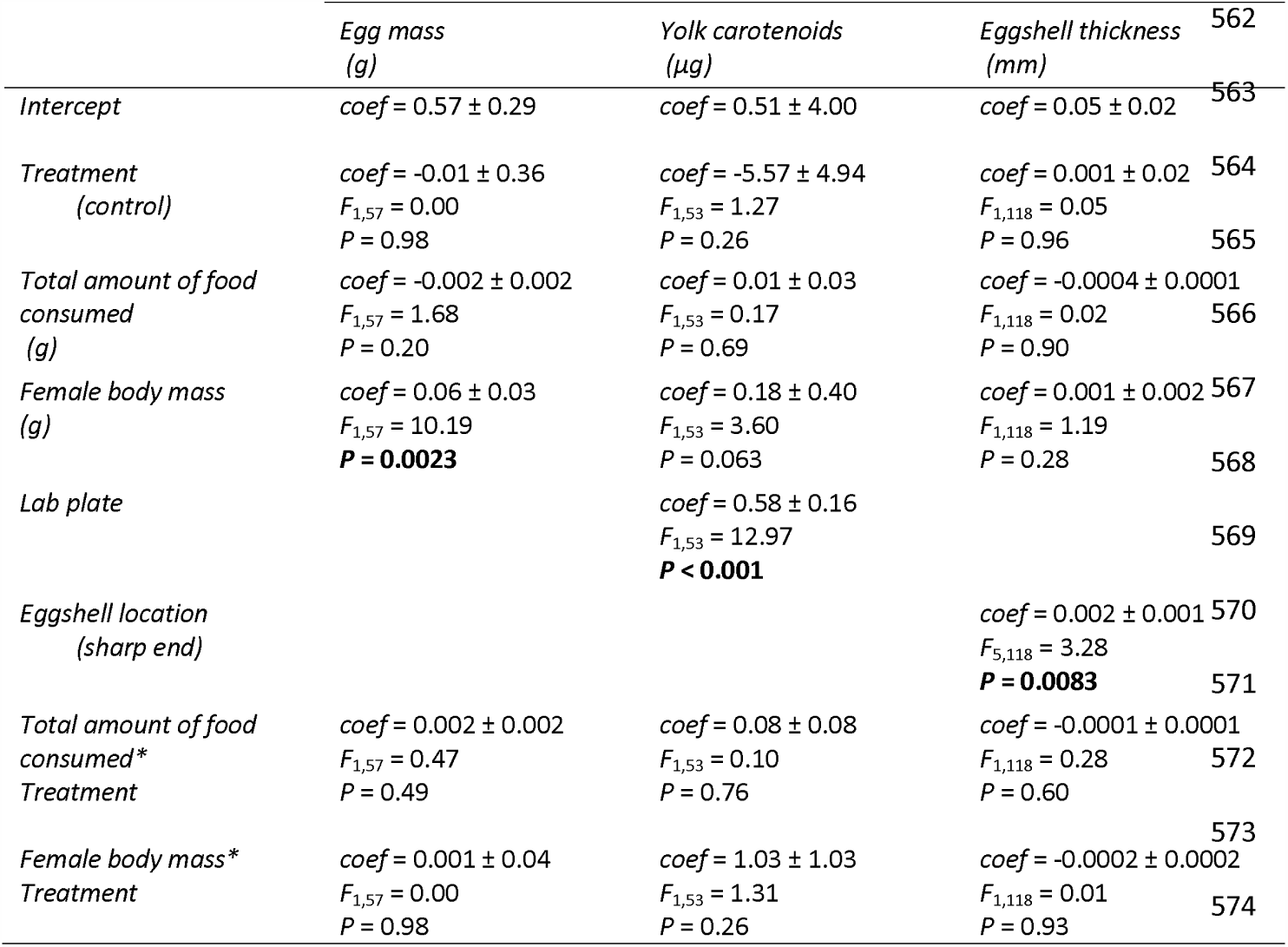
Full models before backward deletion of non-significant interactions showing the effects of treatment on egg quality. General lineal models were performed for egg mass and yolk carotenoid content. A mixed model was performed for eggshell thickness. Coefficients are shown for control nests and sharp end location. Significant differences are marked in bold.

|  | *Egg mass*<br>*(g)* | *Yolk carotenoids*<br>*(μg)* | *Eggshell thickness*<br>*(mm)* | 562 |
| --- | --- | --- | --- | --- |
| *Intercept* | *coef* = 0.57 ± 0.29 | *coef* = 0.51 ± 4.00 | *coef* = 0.05 ± 0.02 | 563 |
| *Treatment*<br>*(control)* | *coef* = -0.01 ± 0.36<br>*F*<sub>1,57</sub> = 0.00<br>*P* = 0.98 | *coef* = -5.57 ± 4.94<br>*F*<sub>1,53</sub> = 1.27<br>*P* = 0.26 | *coef* = 0.001 ± 0.02<br>*F*<sub>1,118</sub> = 0.05<br>*P* = 0.96 | 564<br>565 |
| *Total amount of food*<br>*consumed*<br>*(g)* | *coef* = -0.002 ± 0.002<br>*F*<sub>1,57</sub> = 1.68<br>*P* = 0.20 | *coef* = 0.01 ± 0.03<br>*F*<sub>1,53</sub> = 0.17<br>*P* = 0.69 | *coef* = -0.0004 ± 0.0001<br>*F*<sub>1,118</sub> = 0.02<br>*P* = 0.90 | 566 |
| *Female body mass*<br>*(g)* | *coef* = 0.06 ± 0.03<br>*F*<sub>1,57</sub> = 10.19<br>***P* = 0.0023** | *coef* = 0.18 ± 0.40<br>*F*<sub>1,53</sub> = 3.60<br>*P* = 0.063 | *coef* = 0.001 ± 0.002<br>*F*<sub>1,118</sub> = 1.19<br>*P* = 0.28 | 567<br>568 |
| *Lab plate* |  | *coef* = 0.58 ± 0.16<br>*F*<sub>1,53</sub> = 12.97<br>***P* < 0.001** |  | 569 |
| *Eggshell location*<br>*(sharp end)* |  |  | *coef* = 0.002 ± 0.001<br>*F*<sub>5,118</sub> = 3.28<br>***P* = 0.0083** | 570<br>571 |
| *Total amount of food*<br>*consumed\**<br>*Treatment* | *coef* = 0.002 ± 0.002<br>*F*<sub>1,57</sub> = 0.47<br>*P* = 0.49 | *coef* = 0.08 ± 0.08<br>*F*<sub>1,53</sub> = 0.10<br>*P* = 0.76 | *coef* = -0.0001 ± 0.0001<br>*F*<sub>1,118</sub> = 0.28<br>*P* = 0.60 | 572 |
| *Female body mass\**<br>*Treatment* | *coef* = 0.001 ± 0.04<br>*F*<sub>1,57</sub> = 0.00<br>*P* = 0.98 | *coef* = 1.03 ± 1.03<br>*F*<sub>1,53</sub> = 1.31<br>*P* = 0.26 | *coef* = -0.0002 ± 0.0002<br>*F*<sub>1,118</sub> = 0.01<br>*P* = 0.93 | 573<br>574 |

The yolk carotenoid content of the fifth egg was not affected by treatment (coef. = −0.005 ± 0.25, *F*_1,55_ = 0.02, *P* = 0.98) or by the total amount of food consumed (coef. = 0.00 5 ± 0.01, *F*_1,55_ = 0.37, *P* = 0.72). Females with higher body mass laid eggs with higher yolk carotenoid content (coef. = 0.56 ± 0.23, *F*_1,55_ = 12.15, *P* = 0.019). All other variables and the interactions with treatment had no significant effect (Table 2).

Lutein-supplemented females laid eggs with thinner egg shells than control females (coef. = 0.003 ± 0.001, *F*_1,119_ = 11.68, *P* < 0.001). Eggshell thickness did not vary with female body mass or with the total amount of food consumed (both P > 0.2). Eggshell thickness depended on the eggshell location (coef. = 0.002 ± 0.001, *F*_1,119_ = 3.27, *P* = 0.0084; see also Table 2), sharp end locations being thicker than blunt end locations (least square difference: coef. = 0.002 ± 0.001, *t*_127_ = 2.19, *P* = 0.030), but not to equator locations (coef. = 0.00 1 ± 0.001, *t*_127_ = 1.58, *P* = 0.1163).

## Discussion

We hypothesized that dietary carotenoid availability during egg laying is likely of central importance for female blue tits. Indeed, experimentally enhanced carotenoid availability allowed lutein-supplemented females to have less laying interruptions. However, this came at a potential cost as lutein-supplemented females laid thinner eggs than control females. Intriguingly, other aspects of a female’s laying capacity or the allocation of carotenoids to the yolk were not affected. The potential causes of these findings are discussed below.

### Egg laying capacity

Lutein-supplemented females started eating from the feeders before control females, which was unexpected. This lower food neophobia was independent of female body mass, indicating that it is not driven by female quality differences between groups. One possible explanation could be that blue tits are able to forage selectively on carotenoid-rich foods, via the perception of differences in food colour or smell. Thus, carotenoid treatment could have been more attractive for females, which is supported by a study in great tits (*Parus major*), a species closely related to the blue tit, which chooses to forage on carotenoid-enriched food when exposed to choice tests (Senar et al. 2010). However, even though lutein-supplemented females tended to consume more extra food, this was not reflected in their egg mass (see below).

As expected, we did find an effect of treatment on the number of laying interruptions. Lutein-supplemented females completed their clutch in fewer days than control females, while clutch size was similar. This is the first evidence of the effects of lutein availability on laying interruptions. This effect remained after controlling for female body mass. Previous studies in the blue tit showed that laying interruptions occur frequently in non-food-supplemented females or as a consequence of environmental constraints (Yom-Tov and Wright 1993). Our study focuses on one micronutrient and the results suggest that carotenoids are an essential resource that modulates laying intervals. If blue tit females maintain their clutch size (Nilsson and Svensson 1993; García-Navas and Sanz 2011), but need more than one day for a certain egg, this will ultimately cause a delay in the timing of hatching. Longer laying times can have serious consequences for parental rearing capacity (Perrins 1970), egg viability (Milonoff 1989), and nestling survival (Martin and Hannon 1987; Nilsson 1990; Hochachka 1990). The latter may arise via increased hatching asynchrony, if incubation starts before the clutch is completed, which disadvantages later hatching chicks (Magrath 1990).

Lutein treatment had no effects on laying date, while several studies have shown a positive effect of food supplementation on the timing of reproduction (Martin 1987; Meijer and Drent 1998; Robb 2008). Yet, these effects may relate to caloric restrictions and not to the availability of specific nutrients. However, due to our experimental design we did not expect large effects on laying date. In the current study, feeding both control and lutein-supplemented females with bird fat allowed us to avoid metabolic or energetic effects not related with female carotenoid requirements, and to focus on the role of carotenoids. Furthermore, the onset of feeding may have been too short to affect the onset of laying.

There was no effect on clutch size either (Harrison et al. 2010), despite the fact that previous studies showed that carotenoids may be limiting for egg production (Biard et al. 2005). In the study year, the average clutch size in control and experimental females was 9.36 ± 0.15 (range: 6-14, n=92), which does not differ from the following two years in which females were not supplemented (9.56 ± 0.14, range 4-15, n=206). Thus, it seems that the plasticity in clutch size is limited at least in the study population, and does not depend greatly in food availability or in other specific substances (see also Moreno et al. 1989 in the pied flycatcher *Ficedula hypoleuca*).

### Egg quality

Unexpectedly, we did not find differences in yolk carotenoid content between our treatments. This is in contrast to previous studies showing effects of carotenoid-supplementation on yolk carotenoid content in wild blue tits (Blount et al. 2002; Biard et al. 2005) and in other captive bird species (Surai and Speake 1998; Surai and Sparks 2001; Bortolotti et al. 2003). However, these studies substantially manipulated carotenoids potentially various magnitude orders above the natural consumption (more than 100 times the daily consumption) (daily amount of lutein supplemented: 500 mg in Biard et al. 2005; 1.75 in Remeš et al. 2007). Here, we supplied females with carotenoids within the biological range (0.4 mg in this study; see Partali et al. 1987). Our results suggest that perhaps females used the extra carotenoids for other physiological functions related to self-maintenance and to enhance the laying capacity (Hargitai et al. 2006; Navara et al. 2006; Isaksson et al. 2018). Thus, in the context of a trade-off between allocation to eggs and to self-maintenance (Morales et al. 2008; Giordano et al. 2014), self-maintenance is prioritized until carotenoid supplementation goes beyond the levels required by the female, when it may indeed be reflected in higher yolk carotenoid contents. However, it has to be considered that we assume that carotenoid transfer to the follicle occurs each 24 hours (Salvante and Williams 2002), and, thus, if females had allocated more carotenoids to the yolk, we should have detected differences between treatments in the fifth collected egg. Thus, the lack of an effect on yolk carotenoid content in our study is more likely explained by females using carotenoids for self-maintenance functions than being an artefact of our methodology.

We did not find effects of treatment on egg mass, but both control and lutein-supplemented females received additional resources that could have been used for yolk formation. Yet, egg mass was still positively related to female body mass, which may not necessarily reflect condition differences, but could be due to morphological constraints with larger females being able to lay larger eggs.

We found a treatment effect on eggshell thickness, with lutein-supplemented females laying thinner eggs than control females. One explanation is that laying interruptions allow females to accumulate calcium resources, which is reflected in increased shell thickness. Thus, carotenoid supplementation facilitated egg laying, but less number of laying interruptions during egg sequence prevented the deposition of calcium in the eggshell, either because lutein-supplemented females had less time for foraging on calcium-rich resources between eggs (note that even short foraging bouts result in increased quantities of calcium; Flint et al., 1998) or because their eggs stayed on average less time in the oviduct compared to control eggs. Eggshell is the physical barrier between the embryo and the environment and its thickness has profound consequences on fitness related to incubation efficiency by heat transference (Soliman et al. 1994), microbial infection (D’Alba et al. 2014), water loss (Drent and Woldendorp 1989) and egg viability (Mellanby 1992). Therefore, our results suggest that a consequence effect of laying faster could be a lower accumulation of calcium in the eggshell, which indicates that the combined availability of different resource determines egg quality.

To conclude, this study provides the first evidence that experimentally enhanced carotenoid availability allowed blue tit females to complete their clutch faster. This suggests that carotenoids are a limiting resource in the blue tit, a species that lays large clutches in a very short time interval. Yet, lutein supplementation did not lead to higher yolk carotenoid content, which suggests that females used the extra carotenoids for self-maintenance or to enhance their laying capacities. The supplementation of a single compound, here lutein, also revealed a trade-off between laying in short sequence and calcium deposition in the eggshell, since lutein-supplemented females laid eggs with thinner shells. To summarize, our results emphasize the limiting role that carotenoids play for blue tit females during the egg production.

## Acknowledgments

We thank Lucía Arregui for setting-up the technique for carotenoid analyses.

## Author Contributions

JM, WM and JGC conceived and designed the experiments. JGC, JM and SG performed the experiments. EG performed laboratory analyses. JGC, JM, WM analyzed the data. JGC, JM, WM wrote the manuscript.

## Legends to figures

**Figure 1.**
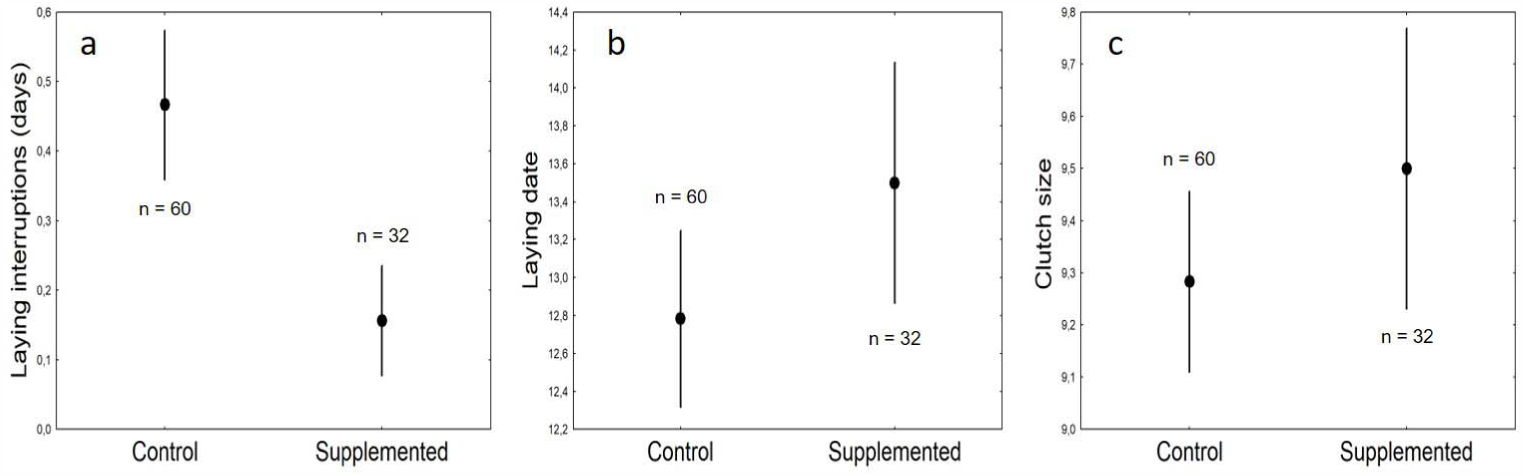
Laying capacity of control and lutein-supplemented blue tit females (*Cyanistes caeruleus*): a) Number of laying interruptions during egg laying (days; GLZ, *P* = 0.033); b) Laying date according to the Julian calendar (GLM, *P* = 0.14); c) Clutch size (GLZ, *P* = 0.75). Error bars denote standard errors (mean ± SE; n=92). Sample sizes for each treatment are shown.

**Figure 2.**
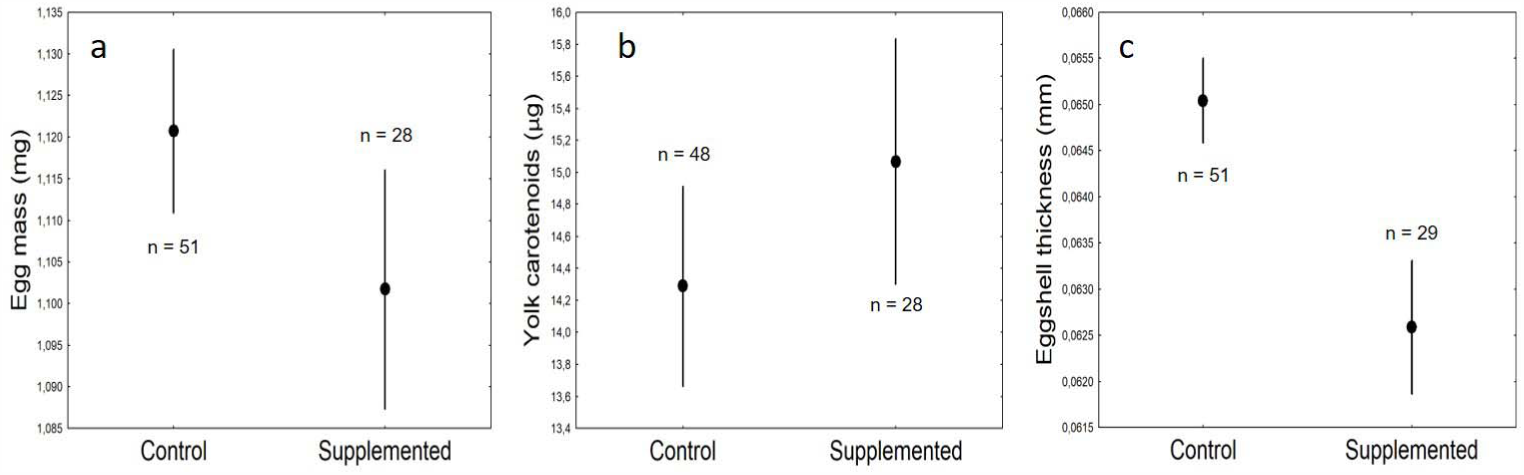
Egg quality measures (mean ± SE) of control and lutein-supplemented blue tit females (*Cyanistes caeruleus*): a) Egg mass of the 5^th^ collected egg (g; GLM, n= 79, *P* = 0.39); b) Yolk carotenoid content of the 5^th^ collected (µg; (GLM, n= 76, *P* = 0.98); c) Eggshell thickness of the 5^th^ collected (mm; GLMM, n= 80, *P* < 0.001); values of shell thickness are the mean measured in all locations pooled (blunt end, sharp end and equator). Sample sizes for each treatment are shown.

## References

Alonso-Álvarez C, Bertrand S, Devevey G, Gaillard M, Prost J, Faivre B, Sorci G (2004) An experimental test of the dose-dependent effect of carotenoids and immune activation on sexual signals and antioxidant activity. Am Nat 164:651–659 https://doi.org/10.1086/424971

Ardia DR, Wasson MF, Winkler DW (2006) Individual quality and food availability determine yolk and egg mass and egg composition in tree swallows Tachycineta bicolor. J Avian Biol 37:252–259 https://doi.org/10.1111/j.2006.0908-8857.03624.x

Astheimer LB (1985) Long Laying Intervals: A Possible Mechanism and Its Implications. Auk 102:401–409 https://doi.org/10.2307/4086789

Biard C, Surai PF, Møller AP (2005) Effects of carotenoid availability during laying on reproduction in the blue tit. Oecologia 144:32–44 https://doi.org/10.1007/s00442-005-0048-x

Biard C, Surai PF, Møller AP (2007) An analysis of pre- and post-hatching maternal effects mediated by carotenoids in the blue tit. J Evolution Biol 20:326–339 https://doi.org/10.1111/j.1420-9101.2006.01194.x

Blount JD, Houston DC, Moller AP (2000) Why egg yolk is yellow. Trends Ecol Evol 15:47–49 https://doi.org/10.1016/S0169-5347(99)01774-7

Blount JD, Houston DC, Surai PF, & Møller, AP (2004) Egg-laying capacity is limited by carotenoid pigment availability in wild gulls Larus fuscus. Proc R Soc B 81:1998–2000 https://doi.org/10.1098/rsbl.2003.0104

Blount JD, Surai PF, Nager RG, Houston DC, Møller AP, Trewby ML, Kennedy MW (2002) Carotenoids and egg quality in the lesser black-backed gull Larus fuscus: a supplemental feeding study of maternal effects. Proc R Soc B 269:29–36 https://doi.org/10.1098/rspb.2001.1840

Bortolotti GR, Negro JJ, Surai PF, Prieto P (2003) Carotenoids in Eggs and Plasma of Red-Legged Partridges: Effects of Diet and Reproductive Output. Physiol Biochem Zool 76:367–374 https://doi.org/10.1086/375432

Bureš S, Weidinger K (2003) Sources and timing of calcium intake during reproduc tion in flycatchers. Oecologia 137:634–647 https://doi.org/10.1007/s00442-003-1380-7

Carey C (1996) Avian Energetics and Nutritional Ecology. In: New York: Chapman

Cohen AA, McGraw KJ, Robinson WD (2009) Serum antioxidant levels in wild birds vary in relation to diet, season, life history strategy, and species. Oecologia 161:673–683 https://doi.org/10.1007/s00442-009-1423-9

D’Alba L, Jones DN, Badawy HT, Eliason CM, Shawkey MD (2014) Antimicrobial properties of a nanostructured eggshell from a compost-nesting bird. J Exp Biol 217:1116–1121 https://doi.org/10.1242/jeb.098343

Drent PJ, Woldendorp JW (1989) Acid rain and eggshells. Nature 339:431 https://doi.org/10.1038/339431a0

Durant JM, Hjermann DØ, Anker-Nilssen T, Beaugrand G, Mysterud A, Pettorelli N, Stenseth NC (2005) Timing and abundance as key mechanisms affecting trophic interactions in variable environments. Ecol Lett 8:952–958. https://doi.org/10.1111/j.1461-0248.2005.00798.x

Eeva T, Lehikoinen E (2010) Polluted environment and cold weather induce laying gaps in great tit and pied flycatcher. Oecologia 162:533–539 https://doi.org/10.1007/S00442-009-1468-9

García-Navas V, Sanz JJ (2011) The importance of a main dish: Nestling diet and foraging behaviour in Mediterranean blue tits in relation to prey phenology. Oecologia 165:639–649 https://doi.org/10.1007/s00442-010-1858-z

Gibb JA, Betts MM (1963) Food and Food Supply of Nestling Tits (Paridae) in Breckland Pine. J Anim Ecol 32:489. https://doi.org/10.2307/2605

Giordano M, Groothuis TGG, Tschirren B (2014) Interactions between prenatal maternal effects and posthatching conditions in a wild bird population. Behav Ecol 25:1459–1466 https://doi.org/10.1093/beheco/aru149

Graveland J (1996) Avian eggshell formation in calcium-rich and calcium-poor habitats: Importance of snail shells and anthropogenic calcium sources. Can J Zool 74:1035–1044 https://doi.org/10.1139/z96-115

Haq A-U, Bailey CA, Chinnah A (1996) Effect of β-carotene, canthaxanthin, lutein, and vitamin E on neonatal immunity of chicks when supplemented in the broiler breeder diets. Poultry Sci 75:1092–1097 https://doi.org/10.1007/978-3-642-75110-3_31

Hargitai R, Matus Z, Hegyi G, Michl G, Tóth G, Török J (2006) Antioxidants in the egg yolk of a wild passerine: Differences between breeding seasons. Comp Biochem Phys B 143:145–152 https://doi.org/10.1016/j.cbpb.2005.11.001

Harrison TJE, Smith JA, Martin GR, Chamberlain DE, Bearhop S, Robb GN, Reynolds SJ (2010) Does food supplementation really enhance productivity of breeding birds? Oecologia 164:311–320 https://doi.org/10.1007/s00442-010-1645-x

Hochachka W (1990) Seasonal decline in reproductive performance of song sparrows. Ecology 71:1279–1288 https://doi.org/10.2307/1938265

Hõrak P, Surai PF, Møller AP (2002) Fat-soluble antioxidants in the eggs of great tits Parus major in relation to breeding habitat and laying sequence. Avian Science 2:123–130

Isaksson C, Johansson A, Andersson S (2008) Egg yolk carotenoids in relation to habitat and reproductive investment in the great tit Parus major. Physiol Biochem Zool 81:112–118 https://doi.org/10.1086/522650

Flint PL, Fowler AC, Bottitta GE, Schamber J (1998) Observations of geese foraging for clam shells during spring on the Yukon-Kuskokwim Delta, Alaska. Wilson Bull 110:411–413

Karadas F, Pappas AC, Surai PF, Speake BK (2005) Embryonic development within carotenoid-enriched eggs influences the post-hatch carotenoid status of the chicken Comp Biochem Phys B 141:244–251 https://doi.org/10.1016/j.cbpc.2005.04.001

Korpimäki E, Hakkarainen H (1991) Fluctuating food supply affects the clutch size of Tengmalm’s owl independent of laying date. Oecologia 85:543–552 https://doi.org/10.1007/BF00323767

Lessells ACM, Dingemanse NJ, Both C (2002) Egg weights, egg component weights, and laying gaps in great tits (Parus major) in relation to ambient temperature. Auk 119:1091–1103

Lessells CM, Both C (2002) Egg Weights, Egg Component Weights, and Laying Gaps in Great Tits (Parus major) in Relation to Ambient Temperature. Auk 119:1091–1103 https://doi.org/10.2307/4090236

Magrath RD (1990) Hatching asynchrony in altricial birds. Biol Rev 65:587–622 https://doi.org/10.1111/j.1469-185X.1990.tb01239.x

Martin K, Hannon SJ (1987) Natal philopatry and recruitment of willow ptarmigan i n north central and northwestern Canada. Oecologia 71:518–524 https://doi.org/10.1007/BF00379290

Matthysen E, Adriaensen F, Dhondt AA (2011) Multiple responses to increasing spring temperatures in the breeding cycle of blue and great tits (Cyanistes caeruleus, Parus major). Glob Change Biology 17:1–16 https://doi.org/10.1111/j.1365-2486.2010.02213.x

McGraw KJ, Adkins-Regan E, Parker RS (2005) Maternally derived carotenoid pigments affect offspring survival, sex ratio, and sexual attractiveness in a colorful songbird. Naturwissenschaften 92:375–380 https://doi.org/10.1007/s00114-005-0003-z

McWhinney SLR, Bailey CA, Panigrahy B (1989) Immunoenhancing effect of β-carotene in chicks. Poultry Sci 68:94

Meijer T, Drendt R (2008) Re-examination of the capital and income dichotomy in breeding birds. Ibis 141:399–414 https://doi.org/10.1111/j.1474-919x.1999.tb04409.x

Mellanby K (1992) The DDT Story. In: Br Crop Pr

Milonoff M (1989) Can Nest Predation Limit Clutch Size in Precocial Birds? Oikos 55:424–427 https://doi.org/10.2307/3565604

Monaghan P, Nager RG (1997) Why don’t birds lay more eggs? Trends Ecol Evol 12:270–274 https://doi.org/10.1016/S0169-5347(97)01094-X

Monaghan P, Nager RG, Houston DC (1998) The price of eggs: Increased investment i n egg production reduces the offspring rearing capacity of parents. P Roy Soc B-Biol Sci 265:1731–1735 https://doi.org/10.1098/rspb.1998.0495

Morales J, Ruuskanen S, Laaksonen T, Eeva T, et al. (2013) Variation in eggshell traits between geographically distant populations of pied flycatchers Ficedula hypoleuca. J Avian Biol 44:111–120 https://doi.org/10.1111/j.1600-048X.2012.05782.x

Morales J, Velando A (2013) Signals in family conflicts. Anim Behav 86:11–16 https://doi.org/10.1016/j.anbehav.2013.04.001

Morales J, Velando A, Moreno J (2008) Pigment allocation to eggs decreases plasma antioxidants in a songbird. Behav Ecol Sociobiol 63:227–233 https://doi.org/10.1007/s00265-008-0653-x

Moreno J, Carlson A (1989) Clutch size and the costs of incubation in the pied flycatcher Ficedula hypoleuca. Ornis Scand 20:123–128

Nager RG, Monaghan P, Houston DC (2000) Within-clutch trade-offs between the number and quality of eggs: Experimental manipulations in gulls. Ecology 81:1339–1350 https://doi.org/10.1890/0012-9658(2000)081[1339:WCTOBT]2.0.CO;2

Navara KJ, Badyaev AV, Mendonça MT, Hill GE (2006) Yolk antioxidants vary with male attractiveness and female condition in the house finch (Carpodacus mexicanus). Physiol Biochem Zool 79:1098–1105 https://doi.org/10.1086/507661

Nilsson JÅ (1990) What Determines the Timing and Order of Nest-Leaving in the Mar sh Tit (Parus Palustris)? In: Blondel J., Gosler A., Lebreton JD., McCleery R. (eds) Population Biology of Passerine Birds. NATO ASI Series (Series G: Ecological Sciences), vol 24. Springer, Berlin, Heidelberg.

Nilsson J-A, Svensson E (1993) The Frequency and Timing of Laying Gaps. Ornis Scand 24:122 https://doi.org/10.2307/3676361

Olson VA, Owens IPF (1998) Costly sexual signals: Are carotenoids rare, risky or required? Trends Ecol Evol 13:510–514 https://doi.org/10.1016/S0169-5347(98)01484-0

Partali V, Liaaen-Jensen S, Slagsvold T, Lifjeld JT (1987) Carotenoids in food chain studies—II. The food chain of Parus SPP. Monitored by carotenoid analysis. Comp Biochem Phys B 4:885–888

Pérez-Rodriguez L, Mougeot F, Alonso-Alvarez C, Blas J, Viñuela J, Bortolotti GR (2008) Cell-mediated immune activation rapidly decreases plasma carotenoids but does not affect oxidative stress in red-legged partridges (Alectoris rufa). J Exp Biol 211:2155–2161 https://doi.org/10.1242/jeb.017178

Perrins CM (1970) The Timing of Birds ‘Breeding Seasons. Ibis 112:242–255 https://doi.org/10.1111/j.1474-919X.1970.tb00096.x

Perrins CM, Birkhead TR (1983) Avian Ecology (Tertiary Level Biology). In: Blackie Academic & Professional.

Robb GN, McDonald RA, Chamberlain DE, Bearhop S (2008) Food for thought: Supplementary feeding as a driver of ecological change in avian populations. Front Ecol Environ 6:476–478 https://doi.org/10.1890/060152

Remeš V, Krist M, Bertacche V, Stradi R (2007) Maternal carotenoid supplementation does not affect breeding performance in the Great Tit (Parus major). Funct Ecol 21:776–783 https://doi.org/10.1111/j.1365-2435.2007.01277.x

Salvador A (2016) Herrerillo común – Cyanistes caeruleus. In: Enciclopedia Virtual de los Vertebrados Españoles. Salvador, A., Morales, M. B. (Eds.). Museo Nacional de Ciencias Naturales, Madrid. http://www.vertebradosibericos.org/

Salvante KG, Williams TD (2002) Vitellogenin dynamics during egg-laying: Daily variation, repeatability and relationship with egg size. J Avian Biol 33:391–398 https://doi.org/10.1034/j.1600-048X.2002.02920.x

Senar JC, Møller AP, Ruiz I, Negro JJ, Broggi J, Hohtola E (2010) Specific appetite for carotenoids in a colorful bird. PLoS ONE 5:1–4 https://doi.org/10.1371/journal.pone.0010716

Sheldon BC, Verhulst S (1996) Ecological immunology: Costly parasite defences and trade-offs in evolutionary ecology. Trends Ecol Evol 11:317–321 https://doi.org/10.1016/0169-5347(96)10039-2

Soliman FNK, Rizk RE, Brake J (1994) Relationship between shell porosity, shell thickness, egg weight loss, and embryonic development in Japanese quail eggs. Poultry Sci 73:1607–1611 https://doi.org/10.3382/ps.0731607

Stearns SC (1992) The Evolution of Life Histories. In: Oxford: Oxford University Press.

Stenning M (2018) The Blue Tit. In: T & A D Poyser.

Surai PF, Sparks NHC (2001) Designer eggs: From improvement of egg compositio n to functional food. Trends Food Sci Tech 12:7–16 https://doi.org/10.1016/S0924-2244(01)00048-6

Surai PF, Speake BK (1998) Distribution of carotenoids from the yolk to the tissues of the chick embryo. J Nutr Biochem 9:645–651 https://doi.org/10.1016/S0955-2863(98)00068-0

Surai PF, Speake BK, Sparks NHC (2001) Carotenoids in avian nutrition and embryonic development. 1. Absorption, availability and levels in plasma and egg yolk. J Poult Sci 38:1–27

Tschirren B, Fitze PS, Richner H (2005) Carotenoid-based nestling colouration and parental favouritism in the great tit. Oecologia 143:477–482 https://doi.org/10.1007/s00442-004-1812-z

Vafidis JO, Vaughan IP, Jones TH, Facey RJ, Parry R, Thomas RJ (2016) The effects of supplementary food on the breeding performance of Eurasian reed warblers Acrocephalus scirpaceus; implications for climate change impacts. PLoS ONE 11 https://doi.org/10.1371/journal.pone.0159933

Valcu CM, Scheltema RA, Schweiggert RM, Valcu M, Teltscher K, Walther DM, Ca rle R, Kempenaers B (2019) Life history shapes variation in egg composition in the blue tit Cyanistes caeruleus. Commun Biol 2 https://doi.org/10.1038/s42003-018-0247-8

Visser ME, Lessells CM (2001) The costs of egg production and incubation in great tits (Parus major). Proc R Soc B-Bio Sci 268:1271–1277 https://doi.org/10.1098/rspb.2001.1661

Walker LK, Thorogood R, Karadas F, Raubenheimer D, Kilner RM, Ewen JG (2014) Foraging for carotenoids: do colorful male hihi target carotenoid-rich foods in the wild? Behav Ecol 25:1048–1057 https://doi.org/10.1093/beheco/aru076

Watson H, Salmón P, Isaksson C (2018) Maternally derived yolk antioxidants buffer the developing avian embryo against oxidative stress induced by hyperoxia. J Exp Biol 221 https://doi.org/10.1242/jeb.179465

Williams, TD, Miller M (2003) Individual and resource-dependent variation in ability to lay supranormal clutches in response to egg removal. Auk 120:481–489 https://doi.org/10.1093/auk/120.2.481

Yom-Tov Y, Wright J (1993) Effect of heating nest-boxes on egg laying in the blue tit (Parus caeruleus). Auk 110:95–99 https://doi.org/10.1093/auk/110.1.95

